# Genetically-encoded photosensitizers enable light-controlled polymerization on living neuronal membranes

**DOI:** 10.1101/2022.12.27.521977

**Authors:** Anqi Zhang, Chandan S. Kadur, Charu Ramakrishnan, Zhenan Bao, Karl Deisseroth

**Affiliations:** Department of Chemical Engineering, Stanford University, Stanford, CA 94305, USA; Department of Bioengineering, Stanford University, Stanford, CA 94305, USA; CNC program, School of Medicine, Stanford University, Stanford, CA 94305, USA; Department of Psychiatry and Behavioral Sciences, Stanford University, Stanford, CA 94305, USA; Howard Hughes Medical Institute, Stanford University, Stanford, CA 94305, USA

## Abstract

The ability to record, stimulate, and modify brains of living animals would unlock numerous research opportunities and create potential clinical interventions, but it is difficult to interface with a living neural network without damaging it. We previously reported a novel approach to building neural interfaces, namely: genetically programming cells to build artificial structures to modify the electrical properties of neurons *in situ*, which opens up the possibility of modifying neural circuits in living animals without surgery. However, the spatiotemporal resolution, efficiency, and biocompatibility of this approach were still limited and lacked selectivity on cell membrane. Here, we demonstrate an approach using genetically-targeted photosensitizers to instruct living cells to synthesize functional materials directly on the plasma membrane under the control of light. Polymers synthesized by this approach were selectively deposited on the membrane of targeted live neurons. This platform can be readily extended to incorporate a broad range of light-controlled reactions onto specific cells, which may enable researchers to grow seamless, dynamic interfaces directly in living animals.

## Main

Living neural networks exhibit complex physical and dynamical design properties that are challenging to couple with external interfaces in a non-invasive manner. For example, the human brain contains nearly 100 billion neurons, and over 100 trillion synaptic connections with nanoscale spatial organization, and the fundamental information-propagation signals for neuronal communication (action potentials) operate on the millisecond timescale. As a result of these challenges, existing devices and tools for studying and modulating the brain remain severely under-engineered in terms of spatiotemporal resolution, sensitivity, and specificity, and cannot achieve high-content, specific, and non-invasive integration with the biological system, due in large part to mismatches in size, scalability, and plasticity with the fundamental biological elements, and the inability to target specific cell types^1,2^.

A fundamentally new approach to addressing this size and scale mismatch is to genetically program specific cells within living tissue, endowing the targeted cellular elements (e.g. neurons in the brain) with the intrinsic ability to build artificial structures with desired form and function. We previously took the initial step in this direction, establishing the field of genetically targeted chemical assembly, or GTCA^3^; in the first instantiation of this concept, genetically-targeted neurons were modified to enable the cells to construct and deposit functional polymers *in situ*^3^. Specifically, an ascorbate peroxidase (Apex2^4^) was expressed in neurons to serve as genetically-targeted catalyst, and used to initiate hydrogen peroxide (H_2_O_2_)-enabled oxidative polymerization of either conductive or insulating polymers^3^. Electrophysiological and behavioral analyses were used to confirm that genetically-targeted assembly of functional polymers altered membrane properties in the anticipated manner, and modulated cell type-specific behaviors in living organisms^3^ according to the GTCA design.

Despite this initial success, the first system exhibited two limitations. First, spatiotemporal resolution was limited: both the Apex2-encoding viral vector and diffusible-monomer solutions were delivered by intraparenchymal fluidic injection into the brain, and thus the Apex2/H_2_O_2_ system could only trigger a one-time oxidative reaction terminating with depletion of the injected supply. To better match the complexity and plasticity of biological assembly, and leverage the rapid advances in light-guidance within biological systems driven in large part by optogenetics^5-8^, we here explored a light-controlled approach, using genetically-encoded photosensitizers^9^ that produce reactive oxygen species (ROS) upon illumination to serve as reaction centers for photopolymerization.

Second, in our previous experiments, the Apex2 peroxidases were not strictly targeted to the plasma membrane, and indeed the majority of Apex2 was retained intracellularly^10^. Developing a system to efficiently place the light-responsive reaction centers on the plasma membrane surface will be critical to most further applications of this technological platform for the following reasons: 1) live cells with intact membranes are not typically permeable to most polymer precursors or other biosynthetic materials, such that insufficient membrane-displayed enzymes will lead to low reaction yield; 2) increasing the number of membrane-displayed enzymes catalyzing the polymerization reactions will allow for reduction in the ROS exposure and light intensity parameters needed, and thus improve biocompatibility; and 3) intracellularly expressed photosensitizers can cause oxidative stress to cells and indeed have long been used for targeted cell ablation^9^, such that moving photosensitizing reactions to the exclusively extracellular space will limit adverse effects on native intracellular chemistry and cell viability (indeed, previous reports have also demonstrated that intracellular polymerization reactions are also toxic to cells and can induce rapid apoptosis^11-13^.)

Here, we present proof-of-concept for light-controlled polymerization in GTCA, triggered by genetically encoded photosensitizers localized to the surface membrane of living neurons. A significantly advanced molecular strategy was used to target these photosensitizers specifically onto the plasma membrane of primary neurons with minimal intracellular retention. Demonstrating success of this design, polymers synthesized by this approach were selectively deposited onto the surface membrane of targeted living neurons.

### *In situ* genetically-targeted photopolymerization on the surface of neurons

An overview of our concept is shown in Fig. 1a. We first designed a DNA plasmid backbone for expressing membrane-targeted photosensitizers in primary neurons. This construct was composed of a neuron-specific human Synapsin (hSyn) promoter, followed by sequences encoding an IgK leader sequence to initially direct the protein into the endoplasmic reticulum (ER), FLAG tags for antibody detection, the photosensitizer, a transmembrane (TM) domain as the membrane targeting anchor, 2A self-cleaving peptides, and enhanced yellow fluorescent protein (YFP). The targeted neurons are thereby expected to express membrane-displayed photosensitizer and cytosolic YFP (the photosensitizer locations can be determined by staining with antibodies targeting the FLAG tags). Upon illumination with blue light, membrane-displayed photosensitizers are anticipated to act as reaction centers, facilitating oxidative radical polymerization upon targeted neurons. Because of the low solubility of resulting polymers, these synthesized polymers are expected to be deposited onto the targeted cell membrane.

**Figure 1.**
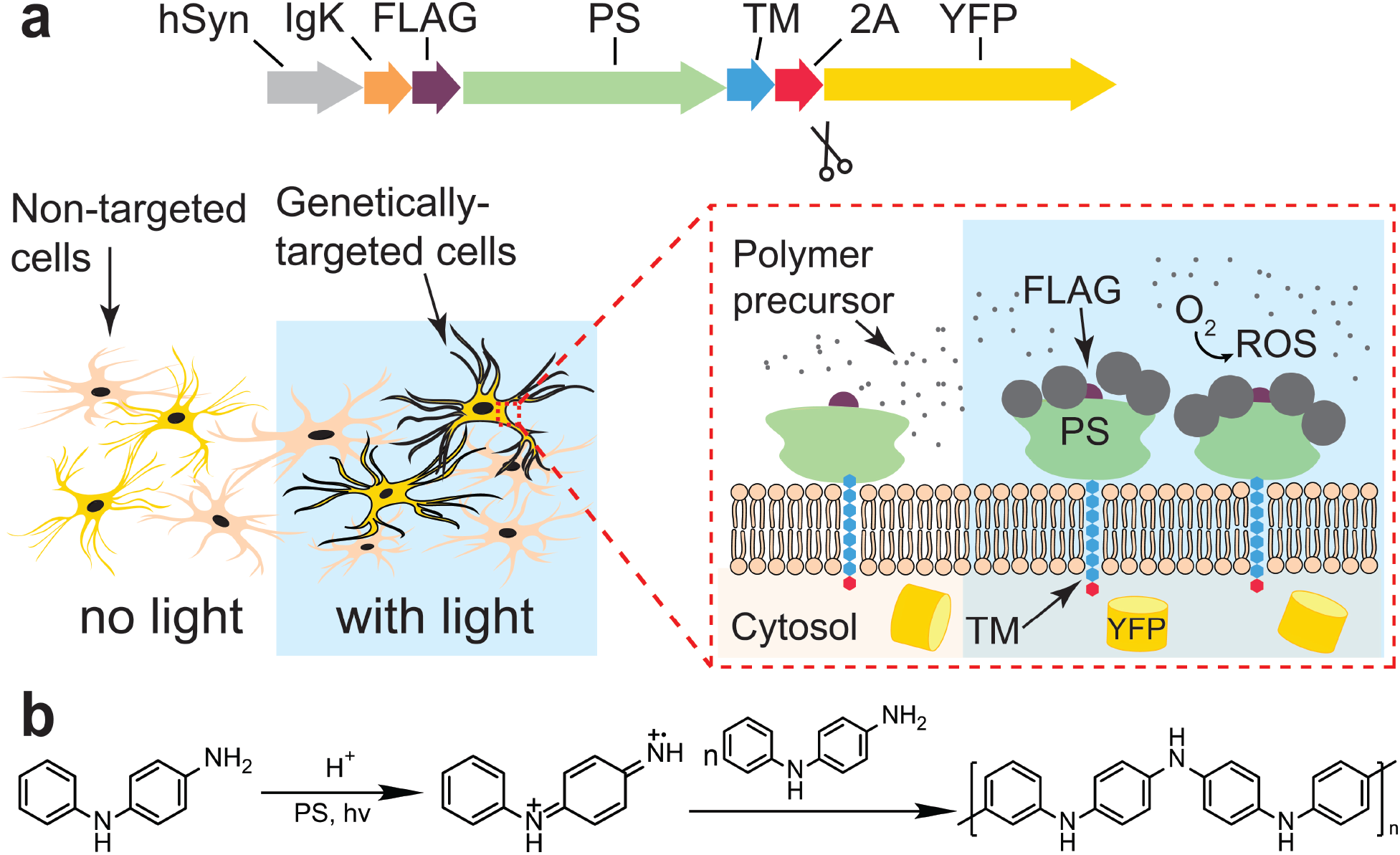
Genetically encoded photosensitizers enabling light-controlled polymerization upon the surface of neurons. (**a**) *Top*, DNA backbone for expressing membrane-displayed photosensitizers (PS). The construct is composed of a neuron-specific hSyn promoter, followed by IgK leader coding sequence, a FLAG tag for antibody detection, the photosensitizer, a transmembrane (TM) domain as the membrane targeting anchor, 2A self-cleaving peptides, and enhanced yellow fluorescent protein (YFP). Thus the targeted cells are expected to express membrane-displayed photosensitizers and cytosolic YFP. *Bottom*, with blue light illumination, photosensitizers expressed on genetically targeted cells are expected to convert oxygen to reactive oxygen species (ROS), thereby catalyzing the oxidative polymerization reaction. Polymer precursors are anticipated to form dark aggregates deposited on the cell surface. Cells in beige depict non-targeted cells. (**b**) Chemical reaction underlying photosensitizer-enabled light-controlled oxidative polymerization of polyaniline from N-phenylenediamine (aniline dimer).

### Selection of photosensitizers

Here, we used N-phenyl-p-phenylenediamine (aniline dimer) for deposition of a conductive polymer, polyaniline (PANI) (Fig. 1b). Critical to the approach described here was our systematic analysis of the genetically-encoded photosensitizers which could be suitable for polymerization on living neurons; **Table 1** lists the genetically-encoded photosensitizers that have been reported so far.

**Table 1.** Genetically-encoded photosensitizer for photopolymerization.

| Protein | MW (kDa) | Ex/Em (nm) | Demonstrated applications | Cell ablation dose | Advantages | Disadvantages |
| --- | --- | --- | --- | --- | --- | --- |
| KillerRed | ~27*2 Dimer<sup>14</sup> | 585/610<sup>14</sup> | Cell ablation<sup>15,16</sup> | 153 J cm<sup>-2</sup> in *C. elegans*<sup>17</sup> | Expression tested in transgenic *C. elegans*, *Drosophila* and zebrafish, and mouse retina with AAV<sup>15,16</sup> | Cannot polymerize DAB<sup>18</sup> |
| miniSOG | ~14 Monomer<sup>19</sup> | 448/528<sup>19</sup> | Cell ablation<br>Polymerization of 3,3 - Diaminobenzidine (DAB)<sup>15,16</sup> | - Cell ablation<sup>20</sup><br>280 J cm<sup>-2</sup><br>- DAB polymerization<br>9.72 J cm<sup>-2</sup> in solution<sup>18</sup><br>120 J cm<sup>-2</sup> in *Drosophila*<sup>21</sup> | Can polymerize DAB, tested in multiple reports<sup>15,16</sup><br>Expression tested in transgenic *C. elegans*, *Drosophila* and zebrafish, and mouse brain<sup>15,16</sup><br>Small size<sup>19</sup><br>Mutants of miniSOG are reported to increase <sup>1</sup>O<sub>2</sub> production by ~10x<sup>18,22-24</sup>. | The short excitation wavelength might cause phototoxicity |
| FAP dL5** | ~25 Monomer<sup>25</sup> | 669/705 | Cell ablation | ~7 J cm<sup>-2</sup><sup>26</sup> | Better tissue penetration<br>Expression tested in transgenic zebrafish<sup>27</sup><br>Efficient energy conversion | Requires 30min-3h incubation in iodine-substituted dye before adding monomer |

The first genetically encoded ROS-generating protein was KillerRed^14^, a phototoxic fluorescent protein derived from a homolog of GFP (anm2CP), which produces ROS upon illumination with red light (excitation maximum of 585 nm). Expression of KillerRed has been tested in transgenic *C. elegans, Drosophila*, zebrafish, and mouse retina^15,16^, and this molecule has been commonly used for cell ablation^16^. While the dimerization tendency of the initial KillerRed largely prevented its use by fusion with proteins of interest, this problem has been addressed with the development of SuperNova, a monomeric version of KillerRed^28^. However, a substantial disadvantage of KillerRed variants is that they cannot polymerize 3,3-Diaminobenzidine (DAB) in EM applications^18^, revealing lower oxidation potential than Apex2.

We systematically considered other possibilities as follows. Efforts to develop genetically-encoded photosensitizers have focused on flavin mononucleotide (FMN)-binding proteins, because FMN is an efficient photosensitizer with a relatively high quantum yield in producing singlet oxygen ^1^O_2_. Mini singlet oxygen generator (miniSOG)^19^ is a fluorescent flavoprotein engineered from Arabidopsis phototropin 2; miniSOG contains 106 amino acids (thus about half the size of Apex2). Notably, illumination of miniSOG generates sufficient ^1^O_2_ to locally catalyze the polymerization of DAB for electron microscopy^15,16^, and new mutants of miniSOG increase ^1^O_2_ production by up to tenfold^18,22-24^. Although miniSOG exhibits fast photobleaching, this effect can be avoided by incorporating other fluorescent proteins for imaging applications^21^. Lastly, a new type of genetically targeted fluorogen-activating protein FAPdL5** was recently developed to generate ^1^O_2_ under near IR (NIR) illumination (669 nm)^25^. Tissue penetration ability of NIR lasers opens up deep tissue applications, but it remains to be tested if the ^1^O_2_ generated by FAPdL5** would be sufficient to trigger polymerization; notably, FAPdL5** also requires 30 min to 3 h incubations in the externally delivered cofactor iodine-substituted dye (MG-2I).

As all three photosensitizers have been expressed on cell membranes, we were able to evaluate ablation efficiency by comparing light doses required for cell ablation (Table 1). Note that in all reports on membrane-targeting photosensitizers, the photosensitizers were expressed on the inner leaflet of the membrane. In our work, the photosensitizers are expressed on the extracellular side, therefore cell ablation dose should be significantly increased, and polymerization on the cell surface will be much easier since ROS do not need to cross the membrane. FAPdL5** requires much lower light levels than KillerRed or miniSOG, indicating that FAPdL5** might be a stronger oxidizing agent for faster polymerization, while both miniSOG and FAPdL5** could be robust general choices for photopolymerization. We decided to first test miniSOG, given its track record of use in oxidative polymerization.

### Membrane localization of miniSOG

Native transmembrane proteins are synthesized in the ER, transported to the Golgi apparatus, and eventually reach the plasma membrane. In designing our construct backbone, we sought to determine if we could use native transmembrane domains as membrane-targeting anchors, beginning by finding domains suitable for recruiting native membrane trafficking machinery in primary neurons. To evaluate membrane expression profiles, we developed antibody staining procedures to compare non-permeabilized and permeabilized cells (Fig. 2a). We found that the transmembrane domain of the T-cell surface glycoprotein CD2^29^ could most efficiently anchor miniSOG on the neuron membrane. In these experiments, miniSOG fused to the FLAG tags was detected with primary antibodies targeting FLAG, which were then labeled with secondary antibodies with Alexa 647 fluorophores. In non-permeabilized cells only membrane anchored miniSOG could be detected, whereas in permeabilized cells both intracellular and extracellular miniSOG were labeled.

**Figure 2.**
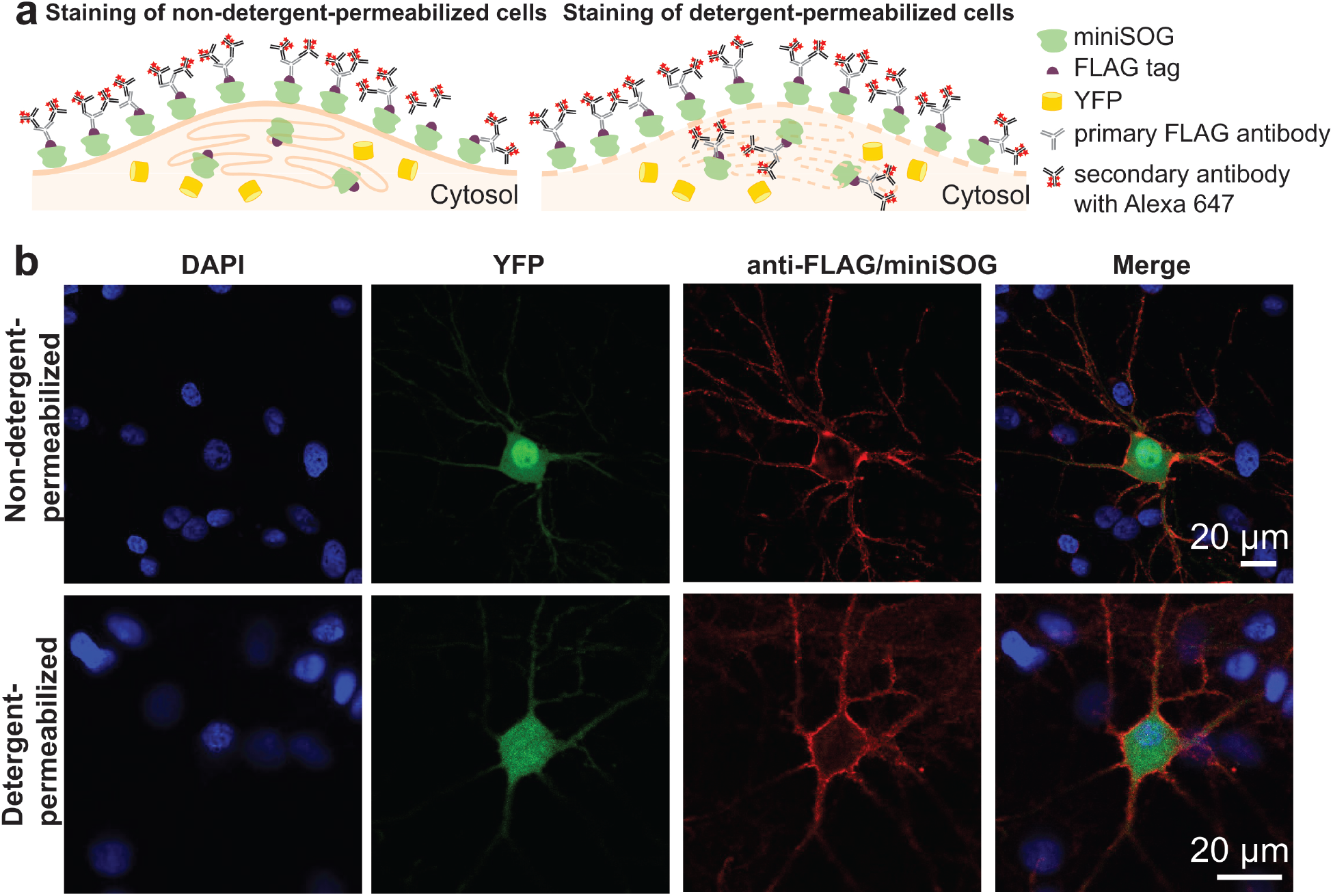
Membrane localization of miniSOG. (**a**) Evaluation of membrane expression by comparing staining results of non-detergent-permeabilized cells and permeabilized cells. miniSOG fused to the FLAG tags was detected with primary antibodies targeting FLAG, which were then labeled via secondary antibodies with Alexa 647 fluorophores. In non-permeabilized cells, only membrane-anchored miniSOG was labeled. In permeabilized cells, both intracellular and extracellular miniSOG were labeled. (**b**) Confocal microscopy of expression and membrane localization of miniSOG in neurons. Seven days post-transfection, cells were fixed, permeabilized or not, and assayed for miniSOG localization in miniSOG(+) cells (cells expressing cytosolic YFP) by immunostaining (cell nuclei were labeled with DAPI as before). The similarity in miniSOG localization in permeabilized and non-permeabilized cells, and the uniform membrane-associated fluorescence, together demonstrated robust membrane localization.

Seven days post-transfection, enzyme localization in miniSOG-CD2 cells (expressing cytosolic YFP) was detected by immunostaining; cell nuclei were labeled with 4’,6-diamidino-2-phenylindole (DAPI). Representative confocal microscopy images (Fig. 2b) showed uniform membrane-associated fluorescence on soma and neurites in the anti-FLAG channel. The similarity of miniSOG localization in permeabilized vs. non-permeabilized cells confirmed the robust membrane-trafficking ability of CD2. We therefore used miniSOG-CD2 neurons as miniSOG(+) cells, with YFP-CD2 neurons as negative control miniSOG(−) cells.

### Photopolymerization on living neurons

Finally, we performed polymerization of PANI on miniSOG(+) and miniSOG(−) neurons (see Methods for reaction conditions) with and without blue light illumination for 10 min (Fig. 3a). Bright field images showed that neurons exhibited dark reaction products as the reactions progressed, and thus reaction progress was represented by changes in brightness of cells relative to background (Fig. 3b). Statistical summary of this brightness decrease revealed robustly and significantly increased polymerization reaction progression on miniSOG(+) neurons under illumination (Fig. 3c).

**Figure 3.**
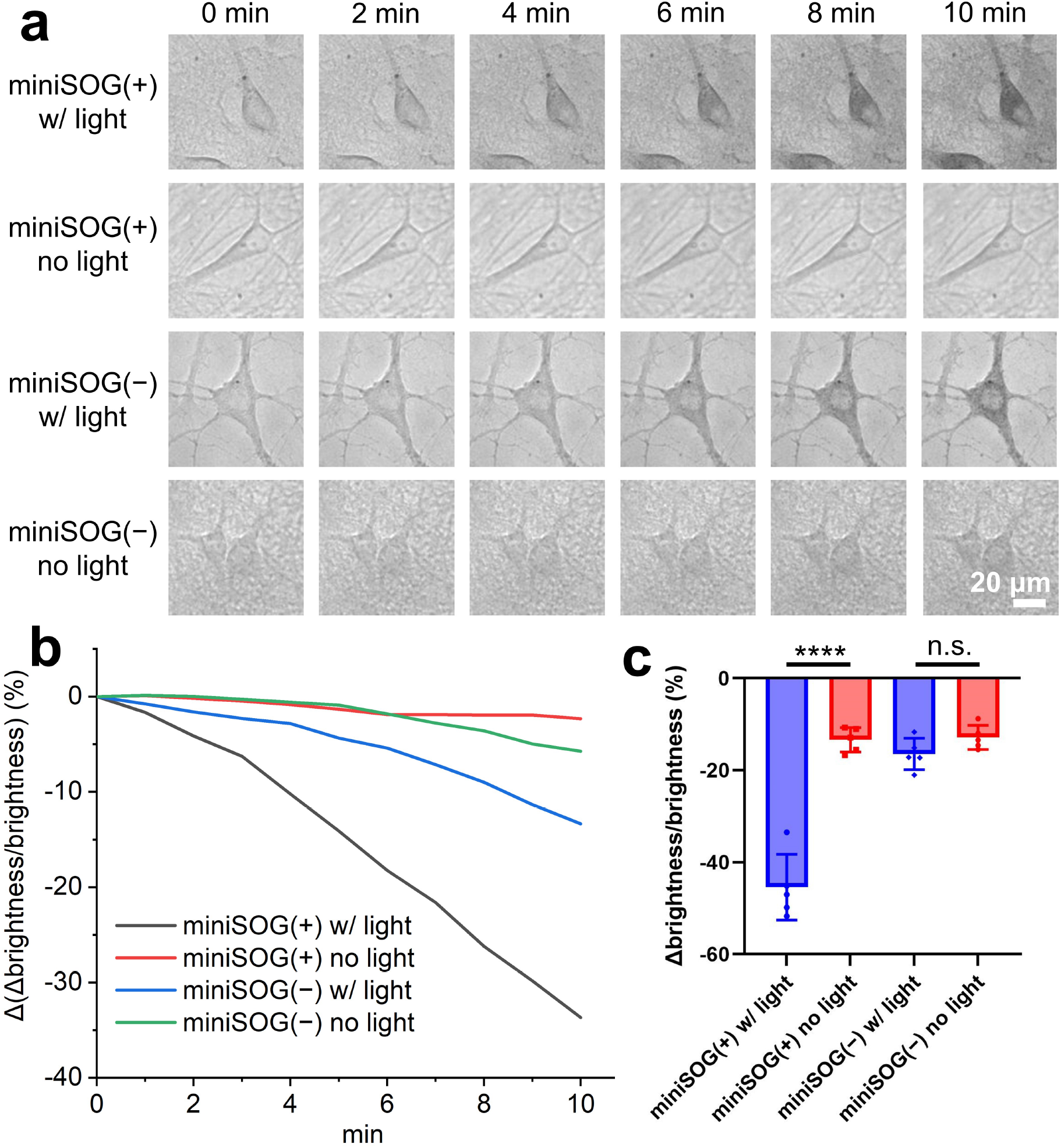
Light controlled polymer deposition. (**a**) Bright field images of live miniSOG(+) and miniSOG(−) neurons with PANI deposition under 20 mW/mm^2^ blue light illumination. (**b**) Representative changes (“Δ”) of the ratio (expressed as %) of brightness differences between neuron and background (“Δbrightness”), to the background brightness, during 10 min polymerization. (**c**) Statistical summary of the brightness ratio after 10 min polymerization. N = 5 cells. Values are means ± s.e.m.; n.s. = nonsignificant, **** = P < 0.0001; two-tailed, unpaired, t-test.

### Conclusion and outlook

This study presents a new GTCA technology for using genetically-targeted photosensitizers to instruct living cells to synthesize functional materials selectively on the plasma membrane. We anticipate that our approach may be readily extended to diverse other areas in the future. For example, a coherent light source can be easily focused to a diffraction-limited spot on the submicron scale, which will allow us to specifically target individual cells and even subcellular structures, perhaps extending ultimately to individual synaptic junctions by leveraging sparse expression strategies for GTCA-components. Additionally, this guided coating of conductive polymers may enable creation of highly refined new conductive pathways between arbitrary cells or structures, thereby letting researchers write new connections into living tissues such as brains. Finally, it is noteworthy that light can be used to initiate a range of photochemical responses (radical chemistry, fluorescence) and photophysical responses (photovoltaic, photothermal) at the illuminated area; a multitude of applications may result in which researchers create atraumatic, highly specific, and scalable interfaces with living organisms.

## Acknowledgements

National Science Foundation Future Manufacturing Program grant (award no. 2037164) supported method development for GTCA on neuronal membranes. Keck Foundation supported miniSOG photo-initiated polymerization. Both grants were awarded to K.D. and Z.B. based on the GTCA concept^3^. Part of this work was performed at the Stanford Nano Shared Facilities (SNSF), supported by the National Science Foundation under award ECCS-2026822.

## Author contributions

A.Z., Z.B., and K.D. and conceived and designed the experiments. A.Z. designed the molecular strategy, developed the polymerization reactions, performed all sample preparation and imaging. C.S.K., C.R., and A.Z. performed neuron culture. A.Z., Z.B., and K.D. analyzed all data, prepared figures and wrote the manuscript with edits from all authors. Z.B., and K.D. supervised all aspects of the work.

### Supplementary note

We note a recent parallel effort derived from GTCA, in which photosensitizers were expressed in neurons but, unlike the case here, not targeted to the cell membrane for performance (Sessler et al., Sci. Adv., 2022, 8, eade1136).

## Methods

### Plasmids construct

The following constructs were designed in SnapGene 5.1.7 and cloned into AAV plasmids with the human Synapsin (hSyn) promoter. All sequences were confirmed with Sanger sequencing (Azenta).

miniSOG(+): hSyn-IgK-FLAGx3-miniSOG-CD2-p2A-t2A-YFP

miniSOG(−): hSyn-IgK-FLAGx3-YFP-CD2-p2A-t2A-YFP

### Neuron culture, transfection, and antibody staining

Primary cultures of postnatal hippocampal rat neurons were prepared on 12-mm coverslips in 24-well plates as described previously^3^. Cells were transfected 6-7 days in vitro (DIV) with various constructs. For each coverslip to be transfected, a DNA-CaCl_2_ mix containing with the following reagents was prepared using calcium phosphate transfection kit (Invitrogen, 44-0052): 1 μg of plasmid DNA, 1 μg of salmon sperm DNA (Invitrogen, 15632-011), 1.875 μL 2M CaCl_2_, and sterile water added for a total volume of 15 μL. Finally, 15 μL of 2X HEPES-buffered saline (HBS) was added, and the resulting 30 μL mix was incubated at room temperature for 20 minutes. The growth medium from each well was removed, saved and replaced with 400 μL pre-warmed minimal essential medium (MEM), and the DNA-CaCl_2_-HBS mix was added dropwise into each well, and the plates were returned to the culture incubator for 60 minutes. Each coverslip was then washed three times with 1 mL of pre-warmed MEM, and placed back to the original neuronal growth medium. Neurons were used 7-8 days post-transfection.

To characterize membrane localization of miniSOG, 7 days post-transfection, cultured neurons expressing different constructs were stained using two protocols, and imaged with Leica TCS SP8 confocal laser scanning microscope:

#### Non-detergent-permeabilized staining

Cultured neurons were washed 3 times with 1 mL of pre-warmed serum free Neurobasal medium (Gibco, 21103049) supplemented with 4% B-27 (Gibco, 17504044) and 2 mM Glutamax (Gibco, 35050061), fixed in 4% paraformaldehyde (PFA) at room temperature for 15 min, and washed 3 times with PBS. Cells were blocked with PBS containing 5% normal goat serum (Jackson ImmunoResearch, 005-000-121) at room temperature for 30 min, and then stained with primary antibody against FLAG (DDDDK tag, Abcam, ab1162) at 1:200 dilution with 5% goat serum at 37 ºC for 1 h, and washed 3 times with PBS. Cells were then stained with Alexa 647 Goat anti-rabbit secondary antibody (Abcam, ab150087) at 1:500 dilution with 5% goat serum at 37 ºC for 1 h, and washed 3 times with PBS. Cells were finally permeabilized in PBS containing 5% goat serum and 0.03% Triton X100 at room temperature for 10 min, and mounted on slides using VECTASHIELD HardSet antifade mounting medium with DAPI (Vector Laboratories).

#### Detergent-permeabilized staining

Cultured neurons were washed and fixed in 4% PFA as described above. Cells were then blocked and permeabilized with PBS containing 5% goat serum and 0.03% Triton X100 at room temperature for 1 h, and then stained with primary antibody and secondary antibody, and mounted with DAPI as described above.

### Light-controlled polymerization reaction

For the polymerization reaction, 3 mM aniline dimer solution was freshly prepared by dissolving 5.5 mg of N-phenyl-p-phenylenediamine (Sigma-Aldrich, 241393) in 10 mL of Tyrode’s solution ∼20 h at room temperature, using a magnetic stir bar. When ready to perform the reaction, the solution was filtered with 0.45 μm syringe filters (Fisher Scientific), and live neurons on coverslips were immersed in the solution, and placed on a Leica CTR6000 inverted microscope equipped with a Lumencor LED Fluorescence Illuminator. The coverslips were illuminated with blue light passed through an excitation filter 434/17 filter and a 40x objective at 20 mW/mm^2^ intensity. Bright field images were taken after every min illumination for 10 min.

### Data analysis

ImageJ software was used for imaging analysis. Statistical analyses for all data were performed with two-tailed unpaired t-test using GraphPad Prism 9. Chemical structures were prepared using ChemDraw 21.0.

